# Mechanism of processive telomerase catalysis revealed by high-resolution optical tweezers

**DOI:** 10.1101/700294

**Authors:** Eric M. Patrick, Joseph Slivka, Bramyn Payne, Matthew J. Comstock, Jens C. Schmidt

## Abstract

Telomere maintenance by telomerase is essential for continuous proliferation of human cells and is vital for the survival of stem cells and 90% of cancer cells. To compensate for telomeric DNA lost during DNA replication, telomerase processively adds GGTTAG repeats to chromosome ends by copying the template region within its RNA subunit. Between repeat additions, the RNA template must be recycled. How telomerase remains associated with substrate DNA during this critical translocation step remains unknown. Using a newly developed single-molecule telomerase activity assay utilizing high-resolution optical tweezers, we demonstrate that stable substrate DNA binding at an anchor site within telomerase facilitates the processive synthesis of telomeric repeats. After release of multiple telomeric repeats from telomerase, we observed folding of product DNA into G-quadruplex structures. Our results provide detailed mechanistic insights into telomerase catalysis, a process of critical importance in aging and cancer.

## Introduction

The ends of human chromosomes, termed telomeres, are composed of G-rich GGTTAG repeats, including a single stranded overhang ^1^. Telomeres shrink by 50 nucleotides with each cell division, due to the end replication problem ^2^. Eventually, short telomeres trigger a cell cycle arrest leading to cell death or senescence, which is an important tumor suppressive mechanism ^1^. A key challenge for telomere maintenance is the G-rich nature of the telomeric repeats in human cells. Specifically, the formation of G-quadruplex (GQ) structures has been shown to affect the replication of telomeres ^3^ and telomerase catalysis ^4^. GQs are very stable nucleic acid structures that are formed by four single stranded telomeric repeats ^5, 6^. To compensate for telomeric DNA lost during DNA replication, continuously proliferating cells such as germ cells, stem cells, and most cancer cells express telomerase ^1^. Telomerase is a reverse transcriptase that processively adds telomeric repeats to the single-stranded overhang at chromosome ends ^7, 8^. Importantly, deficiencies in telomerase lead to premature aging diseases and inappropriate activation of telomerase is a hallmark of 90% of human cancers ^1, 9^. To enhance or inhibit telomerase activity as a therapeutic approach for these diseases, it is critical to determine the molecular mechanisms underlying telomere elongation by telomerase.

Telomerase is composed of the telomerase reverse transcriptase (TERT) protein ^10^, telomerase RNA (TR) ^11^, and several accessory proteins, including the RNA-binding proteins dyskerin and TCAB1 ^12^. The human telomerase ribonucleoprotein (RNP) is organized into two distinct lobes. The catalytic core consists of TERT and the pseudoknot and template regions of TR, while the H/ACA lobe contains the 3’ half of TR bound to dyskerin, TCAB1 and additional H/ACA RNP components^13^. The TERT protein has four domains: the telomerase essential N-terminal (TEN) domain, the telomerase RNA binding domain (TRBD), the reverse transcriptase domain (RT), and C-terminal extension (CTE). The TRBD, RT, and CTE encircle the template region of TR, positioning it in close proximity the catalytic residues in the RT domain ^13, 14^. The TEN-domain directly interacts with the telomeric protein TPP1 to facilitate its recruitment to telomeres ^15–17^, and has been suggested to contribute to processive telomerase catalysis ^18^. While structural and biochemical studies have revealed many molecular details of telomerase catalysis ^8, 13, 14^, how the activities of the various TERT-domains and TR are dynamically coordinated to facilitate processive telomere elongation remains poorly understood.

Cancer cells only contain ~250 assembled telomerase RNPs ^19^ and interactions between telomeres and telomerase are rare ^20^. As a consequence, telomerase is thought to elongate telomeres in a single processive lengthening event ^21, 22^. Consistent with this model, mutations in TERT that specifically reduce telomerase processivity have been shown to have defects in telomere maintenance ^23^. Therefore, telomerase processivity, i.e. the total number of telomeric repeats telomerase synthesizes in a single interaction with the telomere, is the key determinant for the extent of telomere elongation.

To synthesize six nucleotide telomeric repeats, telomerase copies the template region contained in TR ^8^. telomerase processivity requires repeated cycles of melting, translocation, and reannealing of substrate DNA-TR base-pairing ^8^. How telomerase remains associated with substrate DNA, when DNA-RNA base-pairing is disrupted, is unknown. It has been proposed that telomerase contains an anchor site to prevent substrate dissociation during TR translocation ^24–26^, and some reports are consistent with the presence a secondary DNA binding site within telomerase ^18, 27–29^. But there is neither conclusive evidence to demon-strate the existence of an anchor site within telomerase nor mechanistic insight into how such an anchor site would facilitate processive telomerase catalysis.

To address how telomerase processively synthesizes telomeric DNA it is necessary to monitor telomerase catalysis and product release in real time. Bulk primer extension assays are end-point measurements and only report on complete product dissociation, rather than product release while telomerase remains associated with substrate DNA. Previous single-molecule approaches to study telomerase catalysis indirectly detect product formation and have very limited resolution to address product release ^30, 31^. To overcome these challenges, we developed a single-molecule telomerase activity assay utilizing high-resolution optical tweezers. This assay allowed us to directly measure stepwise, processive telomerase activity and simultaneously monitor conformational dynamics of the product DNA. We demonstrate that telomerase tightly associates with its DNA substrate, synthesizing multiple telomeric repeats before releasing them in a single large step. The rate at which product is released from this anchor site closely corresponds to the overall rate of product dissociation from elongating telomerase ^32^, suggesting that it is the main substrate binding site during processive telomere elongation. In addition, we show that product DNA released from telomerase dynamically forms GQs. Analysis of telomeric repeat DNA in the absence of telomerase revealed that GQs rapidly fold and unfold suggesting that they are less static than previously thought. Our results provide detailed mechanistic insight into processive telomerase catalysis, a process critical for telomere maintenance in stem cells and cancer cell survival^1, 9^.

## Results

### A Single-Molecule Telomerase Assay Using High-Resolution Optical Tweezers

To investigate substrate extension by a single telomerase ribonucleoprotein (RNP), we developed a single-molecule telomerase activity assay using dual-trap high-resolution optical tweezers (Fig. 1a). Telomerase and its substrate were attached to separate polystyrene beads (Fig. 1b). The connection between the two beads was formed by the association of telomerase with its substrate DNA (Fig. 1a,b). When applying a low constant force (4.0-4.5 pN) to the tether, substrate elongation by telomerase was measured as an increase in distance between the two beads (Fig. 1a). To attach telomerase to the bead, we utilized a 3xFLAG-HaloTag on TERT, modified with biotin (Fig. 1b,c). Telomerase containing FLAG-TERT and 3xFLAG-HaloTag-TERT modified with biotin, co-purified with similar relative amounts of TR, TCAB1, and dyskerin (Fig. 1c-e), indicating that the tag does not affect telomerase RNP assembly. In addition, the tag had no effect on telomerase activity, stimulation by POT1/TPP1, and the presence of neutravidin did not impact primer extension (Fig. 1f, Supplementary Fig. 1a,b). Together, these results demonstrate that the 3xFLAG-HaloTag modified with biotin fused to the N-terminus TERT does not impact telomerase assembly or catalytic activity, making it well suited to implement our high-resolution optical tweezers-based telomerase assay.

**Fig. 1.**
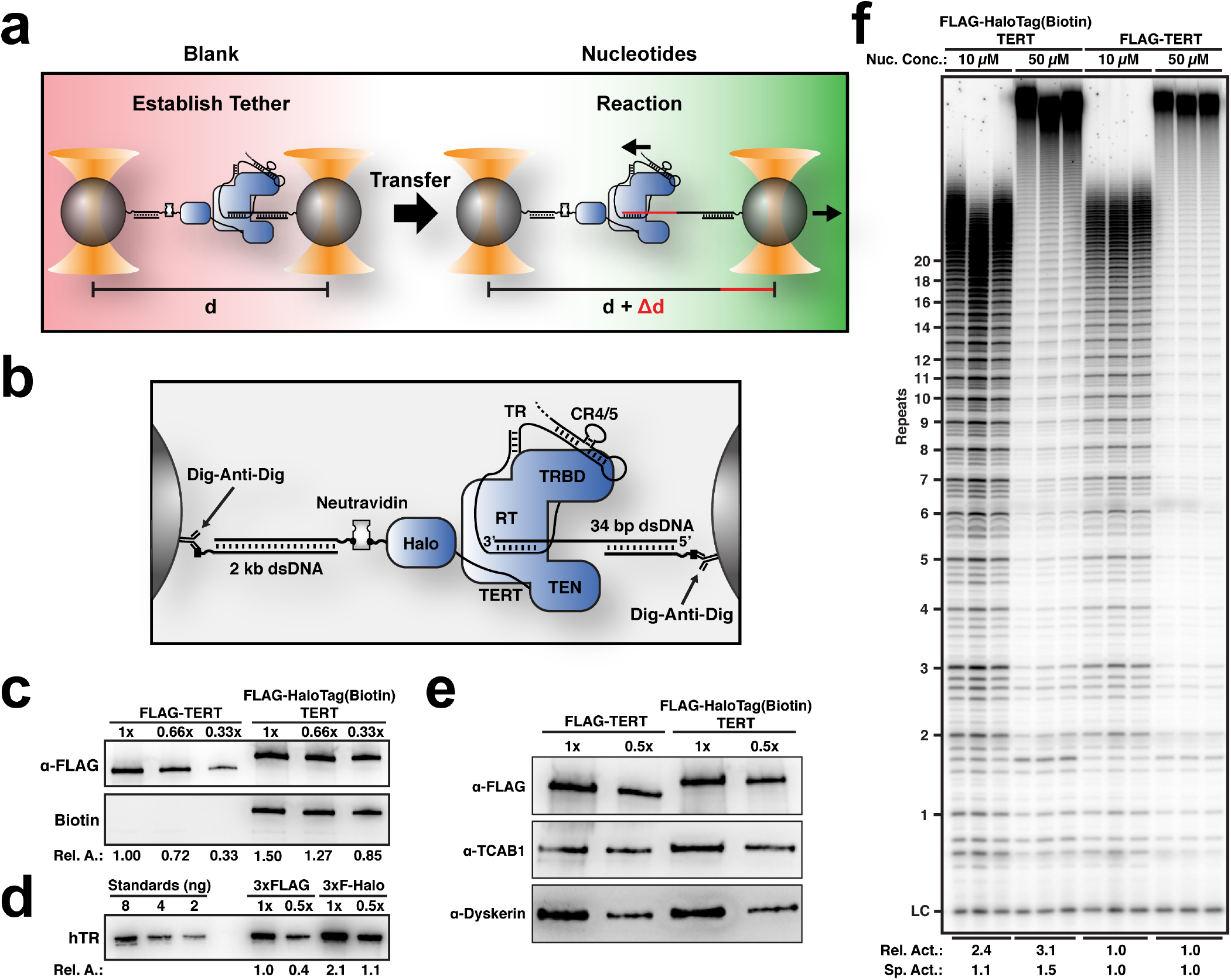
A single-molecule telomerase extension assay using high-resolution dual-trap optical tweezers. **a**, Experimental design to measure processive telomerase catalysis using dual-optical tweezers. **b**, Schematic showing the molecular details of tether formation to monitor DNA elongation by telomerase. **c**, Western blot of purified 3xFLAG- and 3xFLAG-HaloTag-TERT containing telomerase samples, probed with anti-FLAG-M2-HRP (top) and poly-HRP-streptavidin (bottom). **d**, Northern blot of RNA extracted from purified 3xFLAG- and 3xFLAG-HaloTag-TERT containing telomerase samples, probed with three phosphorylated DNA oligonucleotides complementary to TR. Standards are in vitro transcribed full-length TR. **e**, Western blot of purified 3xFLAG- and 3xFLAG-HaloTag-TERT containing telomerase samples, probed with anti-FLAG-M2-HRP (top), TCAB1 (middle), and dyskerin (bottom) antibodies. **f**, Direct telomerase primer extension assay of 3xFLAG-HaloTag-TERT modified with biotin and 3xFLAG-TERT containing telomerase in reaction buffer containing 10 μM or 50 μM dNTPs and 50 mM KCl (LC = loading control). Relative activity is total lane intensity normalized to LC and the respective 3xFLAG-TERT telomerase activity. Specific activity is additionally normalized to the relative amount of TR (Fig. 1d).

### Telomerase Releases Multiple Telomeric Repeats in a Single Step

Using this single-molecule telomerase assay, we set out to analyze how telomerase processively synthe-sizes telomeric repeats. If substrate DNA is only bound to telomerase by base-pairing to TR we would expect to observe a sequence of single nucleotide (nt) addition events as a result of RNA template movement (Supplementary Fig. 2a). In contrast, if the substrate also binds to telomerase at a secondary anchor site delaying nascent DNA strand release, extension events of one or more 6 nt telomeric repeats should be observed (Supplementary Fig. 2b). To monitor telomerase activity, tethers were formed in blank solution lacking nucleotides (Fig. 1a). After assuring that tethers had the expected length and a brief control period, during which no activity was detected (Supplementary Fig. 2c,d), tethers were transferred into the adjacent solution containing nucleotides to initiate telomerase catalysis (Fig. 1a). We observed telomerase activity as a progression of stepwise increases in the tether length (Fig. 2a). The total increase in extension ranged from approximately 20-200 nucleotides (nt, Fig. 2b). The observed elongation rates for the majority (56%) of traces ranged from 8-50 nt per minute (Fig. 2c), which closely agrees with rates measured in bulk experiments ^19, 33^, indicating that the applied force does not alter telomerase activity. The elongation step sizes were broadly distributed, ranging from 5 nt to > 100 nt with the vast majority (86%) being greater than two telomeric repeats (12 nt, Fig. 2d). This implies that telomerase binds to the DNA substrate at a secondary site in addition to the TR-substrate base pairing, allowing multiple telomeric repeats to be synthesized before releasing them in a single large step.

**Fig. 2.**
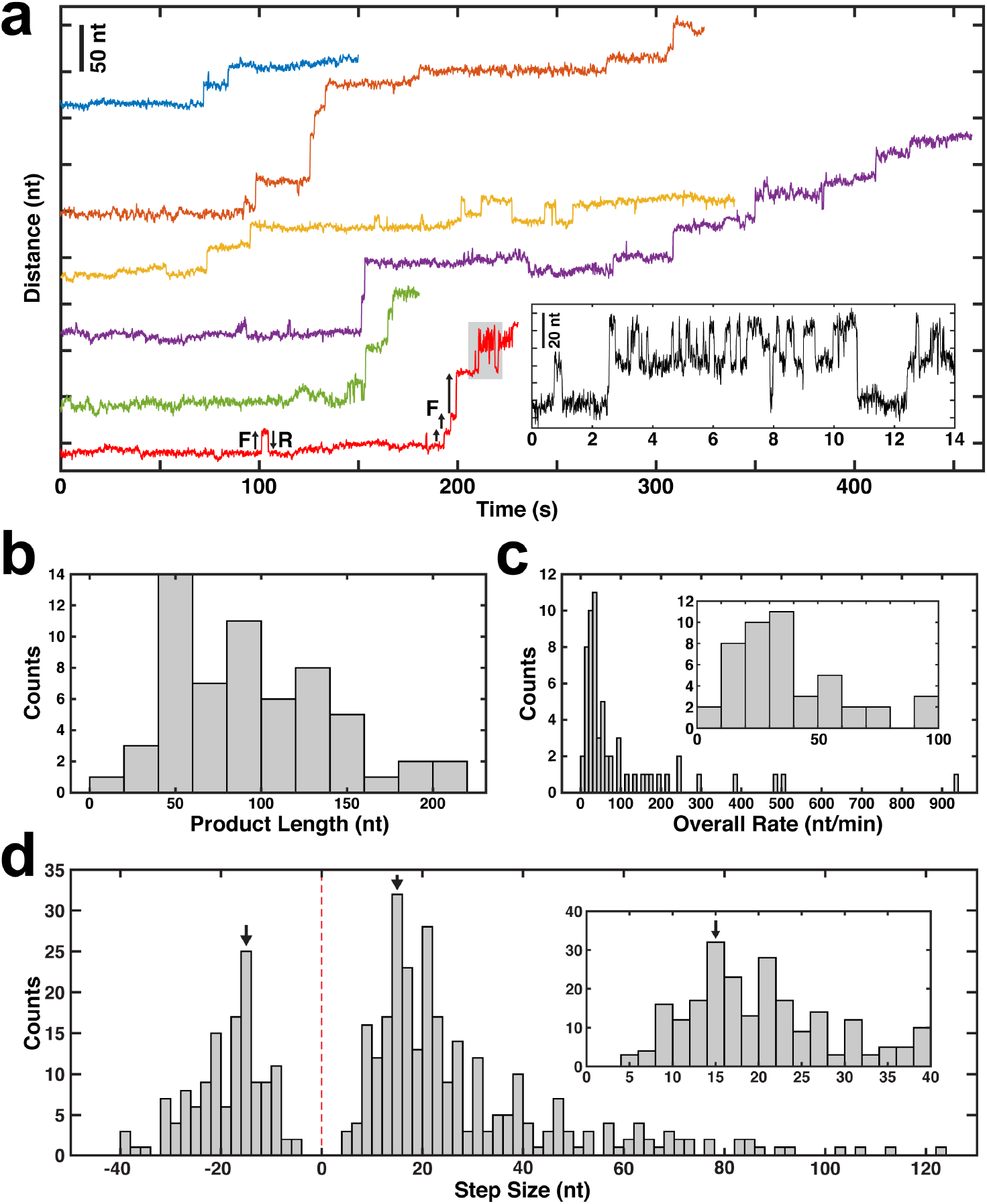
Telomerase releases multiple telomeric repeats in a single step. **a**, Representative time trajectories (120 ms per data point) of telomerase catalysis after transfer into nucleotide containing buffer (t = 0) showing processive elongation, forward (F) and reverse (R) stepping. Inset: Zoom-in of region of bottom-most example trace (in red) corresponding to the region shaded gray. **b**, Distribution of total tether elongation by telomerase (n = 60 traces). **c**, Distribution of overall elongation rate for all telomerase elongation trajectories (n = 60 traces). **d**, Distribution of the observed tether elongation step sizes of telomerase trajectories (n = 60 distinct tethers, 417 steps total). Inset: zoom-in around the forward step distribution peak. Arrows mark a 15 nt magnitude step size which is the expected GQ folding and unfolding step size.

### Telomerase Processively Synthesizes Telomeric Repeats by Stepwise Release of Product DNA

In cancer cells, telomerase is thought to elongate telomeres by 50 nt in a single processive elongation event^22^. Given that the majority of elongation steps (68%) are shorter than 30 nt (Fig. 2d), this would require multiple anchor site release events prior to complete product dissociation. To determine the global progression of substrate elongation by telomerase we analyzed the transition matrix of all traces (Fig. 3a). The first step of all trajectories was a forward step, after which we observed both increases and decreases in tether length (Fig. 3a, Fig. 2a,d). Reverse steps were less frequent than forward steps and, unlike tether extensions, never exceeded 40 nt in length (Fig. 3a, Fig. 2d). Smaller reverse steps frequently matched the preceding forward step and occasionally repetitive forward-reverse step cycling was observed (Fig. 3a, Fig. 2a). On average, telomerase elongated substrates by approximately 72 nucleotides during the first five state transitions (Fig. 3b). After this initial elongation, the average tether length remained constant from step 6 to 10 (Fig. 3b). This lack of further extension coincided with an increased frequency of reverse steps and the appearance of peaks in the step size distribution at approximately 15 nucleotides in both forward and reverse directions (Fig. 3b,c). These observations demonstrate that telomerase can undergo multiple rounds of DNA release from the anchor site without complete product dissociation.

**Fig. 3.**
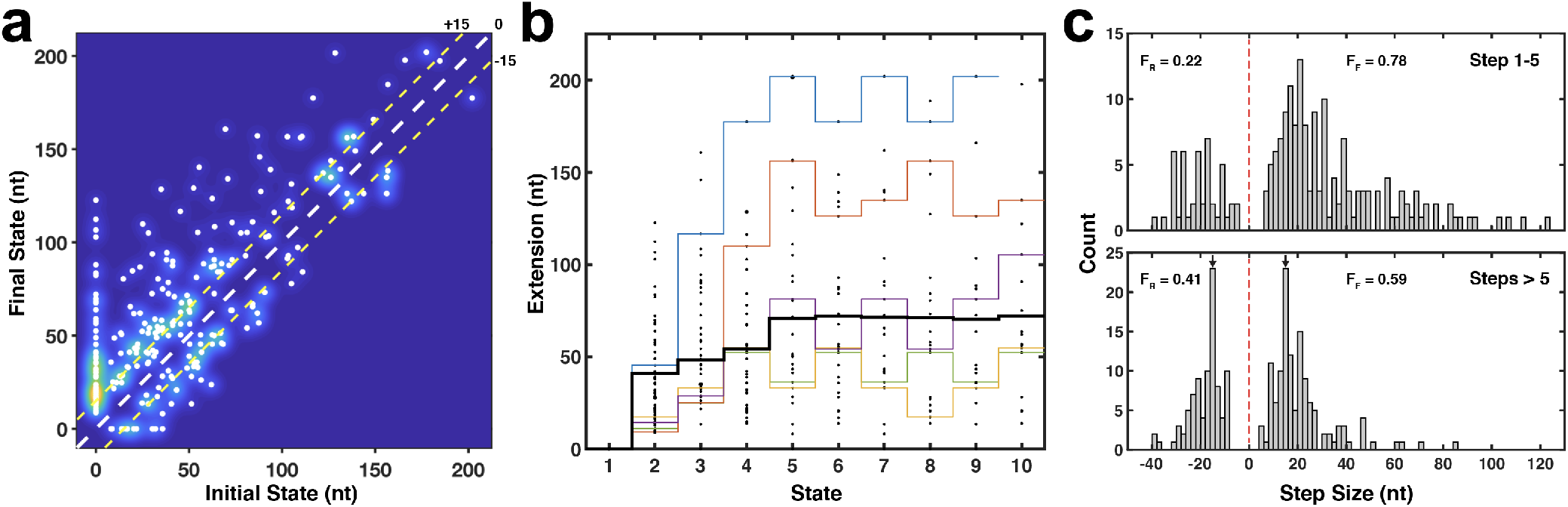
Analysis of processive DNA synthesis and product release by telomerase. **a**, Kernel density plot of final vs. initial states associated with each step in Fig. 2d. The diagonal dashed white line is a guide dividing forward (above line) vs. reverse (below line) steps. The upper and lower dashed yellow lines are guides indicating 15 nt steps forward vs reverse respectively. **b**, Dot plot of the extension for the first 10 states of all telomerase trajectories (n = 60), including the mean extension (black) and individual trajectories (color). **c**, Distribution of step sizes for the first 5 steps (top) and all subsequent steps (bottom) for all trajectories (n = 60 tethers, 417 steps). Arrows mark a 15 nt magnitude step size. F_F_ and F_R_ indicate fractions of forward and reverse steps, respectively.

### Product DNA Released by Telomerase Dynamically Folds into G-Quadruplexes Structures

We next sought to determine the molecular mechanisms that underlie the reverse steps observed in telomerase catalysis trajectories. Reverse steps could be the consequence of two distinct processes: 1) Rebinding of the product to the anchor site, or 2) folding of product into a G-quadruplex (GQ) structure, which has been shown to affect telomerase catalysis ^4^. The most common reverse step size is approximately 15 nt (Fig. 2d, Fig. 3c), which is approximately the expected change in extension for a single GQ folding event under the experimental conditions (Supplementary Fig. 3) ^34^. To test whether GQ folding is responsible for the reverse steps, we conducted telomerase extension experiments in the presence of lithium chloride. Unlike potassium, lithium cations do not support GQ formation ^35^. In the presence of lithium, we observed similar processive telomerase extension, including reverse steps (Fig. 4a). The overall extension rate was slightly reduced, consistent with bulk experiments (Fig. 4b) ^4^. However, lithium eliminated the peaks centered at 15 nt in thestep size distribution in both the forward and reverse directions (Fig. 2d, Fig. 4c,d, arrows), consistent with the interpretation that GQ dynamics are the molecular mechanism underlying these length changes. The remaining reverse steps likely stem from rebinding of the anchor site to product DNA.

**Fig. 4.**
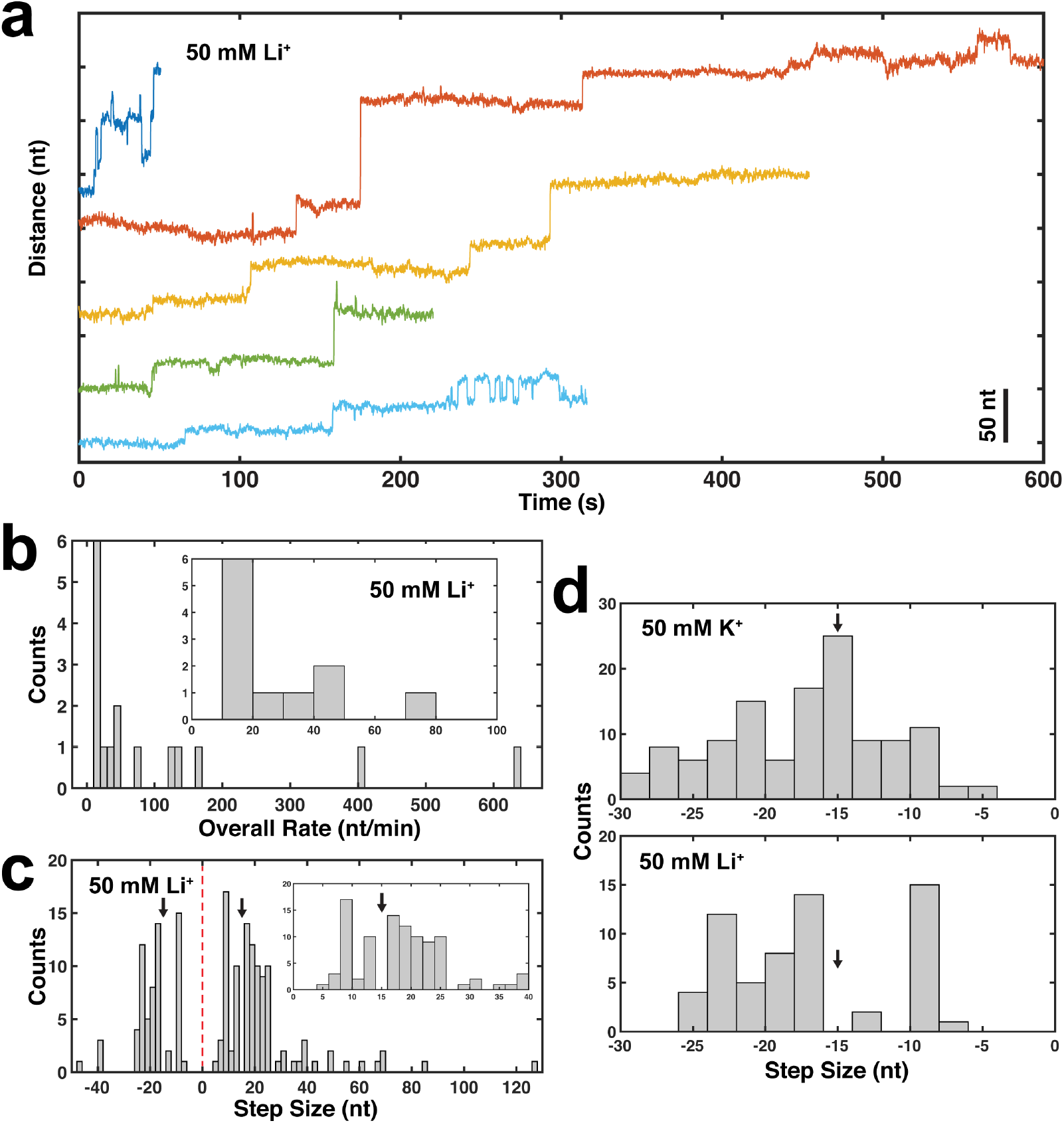
Analysis of telomerase catalysis in the presence of LiCl. **a**, Representative time trajectories (120 ms per data point) of telomerase catalysis in reaction buffer containing 50 mM LiCl after transfer into dNTP containing buffer (t = 0). **b**, Distribution of overall elongation rate in the presence of 50 mM LiCl for all telomerase elongation trajectories (n = 16 traces). **c**, Distribution of the step sizes of telomerase trajectories in reaction buffer containing 50 mM LiCl (n = 16 traces, 171 steps). **d**, Distribution of reverse step sizes of telomerase trajectories in reaction buffer containing 50 mM KCl (n = 60 traces, top) and 50 mM LiCl (n = 16 traces, bottom).

### Telomeric Single Stranded DNA Rapidly Folds and Unfolds G-Quadruplexes Structures

To verify that GQ folding and unfolding occurs with the predicted step size of 15 nt and on time scales relevant for our experiments, we performed tweezers experiments tethering GQ sequences without telomerase. A single stranded DNA (ssDNA) sequence capable of forming one GQ dynamically folded and unfolded into two distinct folded extensions (F1 and F2, with extensions 18.0±0.1 nt and 22.1±0.2 nt, respectively, Fig. 5a,b). These step sizes were consistent with the length changes expected for the formation of different GQ topologies (Supplementary Fig. 3). Simultaneous Förster resonance energy transfer (FRET) measurements ^36^ confirmed the presence of two folded extensions (Fig. 5a, Supplementary Fig. 4a). An extended ssDNA sequence able to form up to three GQ structures dynamically folded and unfolded with step sizes comparable to the single GQ sequence (17.0±0.5 nt and 19.8±0.5 nt, Fig. 5c,d). Both force-extension, and force-feedback measurements confirmed the presence of up to three GQs in the extended telomeric ssDNA sequence (Fig. 5c, Supplementary Fig. 4f). For the single GQ tethers, the dwell time distribution of all folding and unfolding events was fit by the sum of a minimum of three exponential decay functions with a mean dwell time of 11 s (Fig. 5e). The folding dwell time distribution was a single exponential as expected (*τ*_f_=14.2±0.8 s), while unfolding distribution from both folded extensions was fit by a sum of three exponentials, consistent with the formation of at least three distinct GQ structures with varying stabilities (Fig. 5e-g). Further separating the unfolding dwell times by folded extensions F1 and F2 revealed that both folded extensions unfolded with two distinct rate constants (Supplementary Fig. 4b-e). This implies that each folded extension is composed of at least two GQ conformations, yielding a total of at least four total folded states, in agreement with previous observations ^37^. The distribution of all dwell times for the multi-GQ was fit by a minimum of three exponentials with a mean dwell time of 4 s (Supplementary Fig. 4g). Similar to the single GQ, the folding dwell time distribution of multi GQ tethers from the completely unfolded state was a single exponential (τ_f_=3.1±0.3 s) (Supplementary Fig. 4h). In contrast, the unfolding dwell time distribution for multi GQ tethers was fit by a sum of three exponentials, which reflects the complexity of the underlying conformational dynamics (Supplementary Fig. 4i). These results confirm that single stranded telomeric DNA can rapidly fold and unfold GQs under our experimental conditions. In addition, the step sizes of GQ dynamics largely overlapped with the predicted length change for GQ formation.

**Fig. 5.**
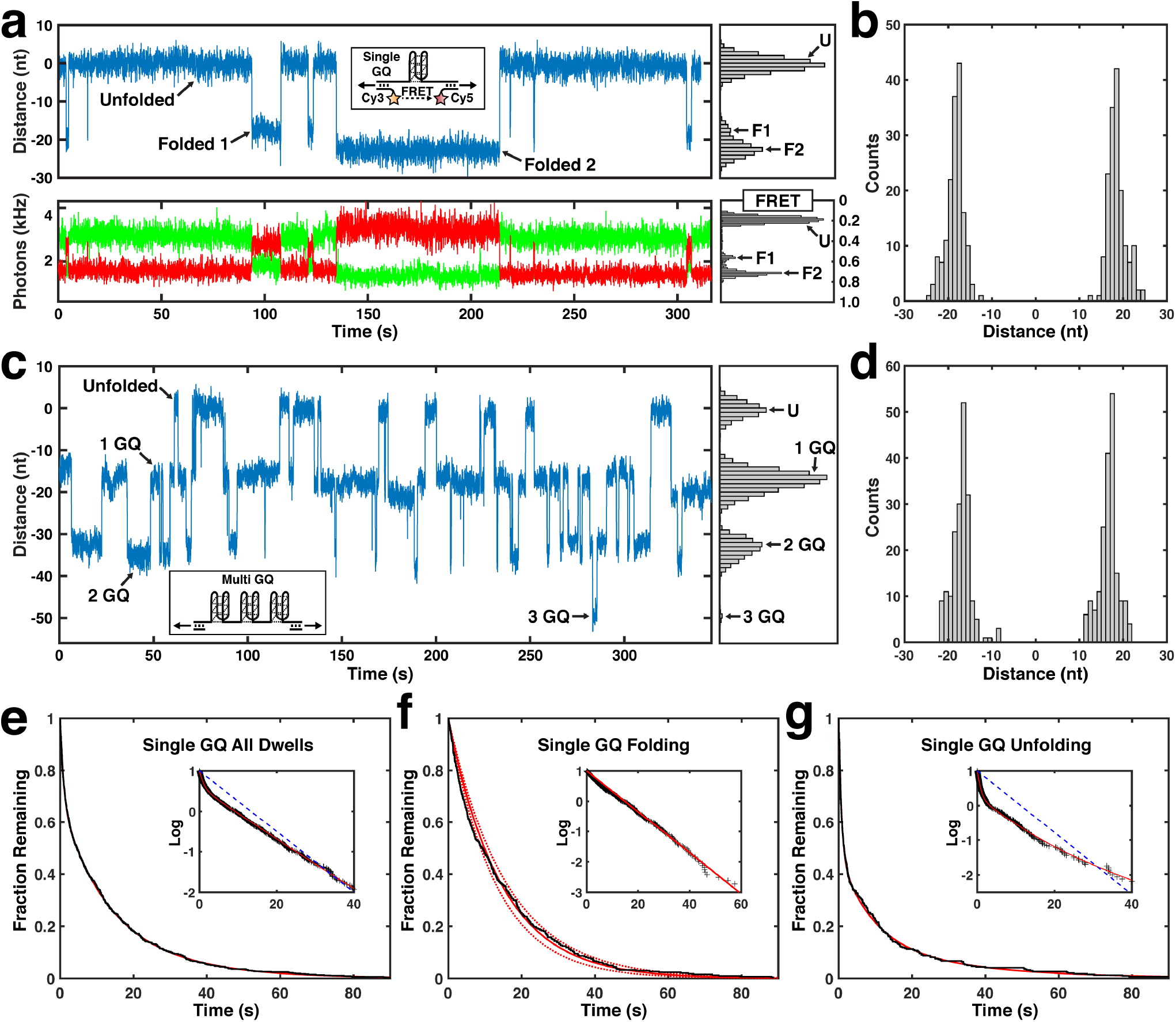
Single stranded telomeric repeat DNA dynamically folds into G-quadruplex structures. **a**, Representative time trajectories and histograms of the folding dynamics of telomeric repeat ssDNA capable of forming a single GQ, showing simultaneous trap distance (top, 48 ms per data point) and FRET measurements (bottom, donor in green, acceptor in red, 48 ms per data point) for a single tether. **b**, Step size distribution of folding and unfolding events observed in single GQ tethers (n = 22 tethers, 344 steps). **c**, Representative time trajectory and histogram of the folding dynamics of telomeric repeat ssDNA capable of forming up to three GQs (48 ms per data point). **d**, Step size distribution of folding and unfolding events observed in multi GQ tethers (n = 6 tethers, 390 steps). **e**, Inverse cumulative distribution of all folding and unfolding dwell times single GQ tethers (black staircase, n = 22 tethers, 701 dwell times). Solid red line indicates fit to the sum of three exponential decays (*τ*_u1_ =0.95±0.22, *τ*_u2_=15±2, and *τ*_u3_=15±3 s, weights of 33±4%, 64±5% and 3±5%). Inset: Same distribution and fit on LOG scale, blue dashed line indicates single exponential decay fit. **f**, Inverse cumulative distribution of dwell times for folding steps of single GQ tethers (black staircase, n = 22 tethers, 351 dwell times). Solid red line indicates a single exponential fit, dashed lines the 90% confidence intervals. Inset: Same distribution and fit on LOG scale. **g**, Inverse cumulative distribution of dwell times for unfolding steps of single GQ tethers (black staircase, n = 22 tethers, 350 dwell times). Solid red line indicates fit to the sum of three exponential decays (*τ*_u1_=0.73±0.12, *τ*_u2_=9.0±3.2, and *τ*_u3_=29±17 s, weights of 52±5%, 32±9% and 15±14%). Inset: Same distribution and fit on LOG scale, blue dashed line indicates single exponential decay fit.

### Telomerase Tightly Binds Substrate DNA at the Anchor Site

The affinity of telomerase for substrate DNA is a key contributing factor to processive telomere repeat synthesis ^38^. To determine the affinity of ssDNA for the telomerase anchor site, we analyzed the dwell times prior to the first elongation step of all telomerase trajectories. Since product release is a pre-requisite for the observation of the initial tether extension, this dwell time should reflect the dissociation rate of ssDNA from the telomerase anchor site. Consistent with this hypothesis, the dwell time distribution prior to the first step was well fit by a single exponential decay with a time constant of 97±12 s (Fig. 6a). After the initial extension step, telomerase could exist in at least two configurations: telomerase RNPs with released ssDNA that is either unfolded or folded into a GQ (Fig. 7). Indeed, the dwell time distributions for the second and third steps fit to the sum of two exponentials (Fig. 6b,c). The slow time constants for the second and third steps (77±20 s, 61±13 s, weights 38±9%, 30±9%) were comparable to the anchor site release time. The fast time constants were approximately 30-fold faster (2.1±0.5 s, 1.4±0.3 s, weights 62±8%, 70±9%), indicating that a second, more rapid, process is occurring in addition to anchor site release. Analysis of the dwell times across the first 10 transitions of all trajectories revealed a gradual decrease of the mean dwell time from approximately 95 s to 10 s from steps 1 to 5 (Fig. 6d). The mean dwell time remained constant from steps 5 to 10 (Fig. 6d). The reduction in dwell time to approximately 10 s is concomitant with the increased frequency of reverse steps (Fig. 3b,c) and is very close to the mean dwell time of 11 s that we measured for the single GQ sequence folding and unfolding (Fig. 5e). These observations suggest that GQ folding and unfolding is the molecular process underlying the faster rate in tether length changes.

**Fig. 6.**
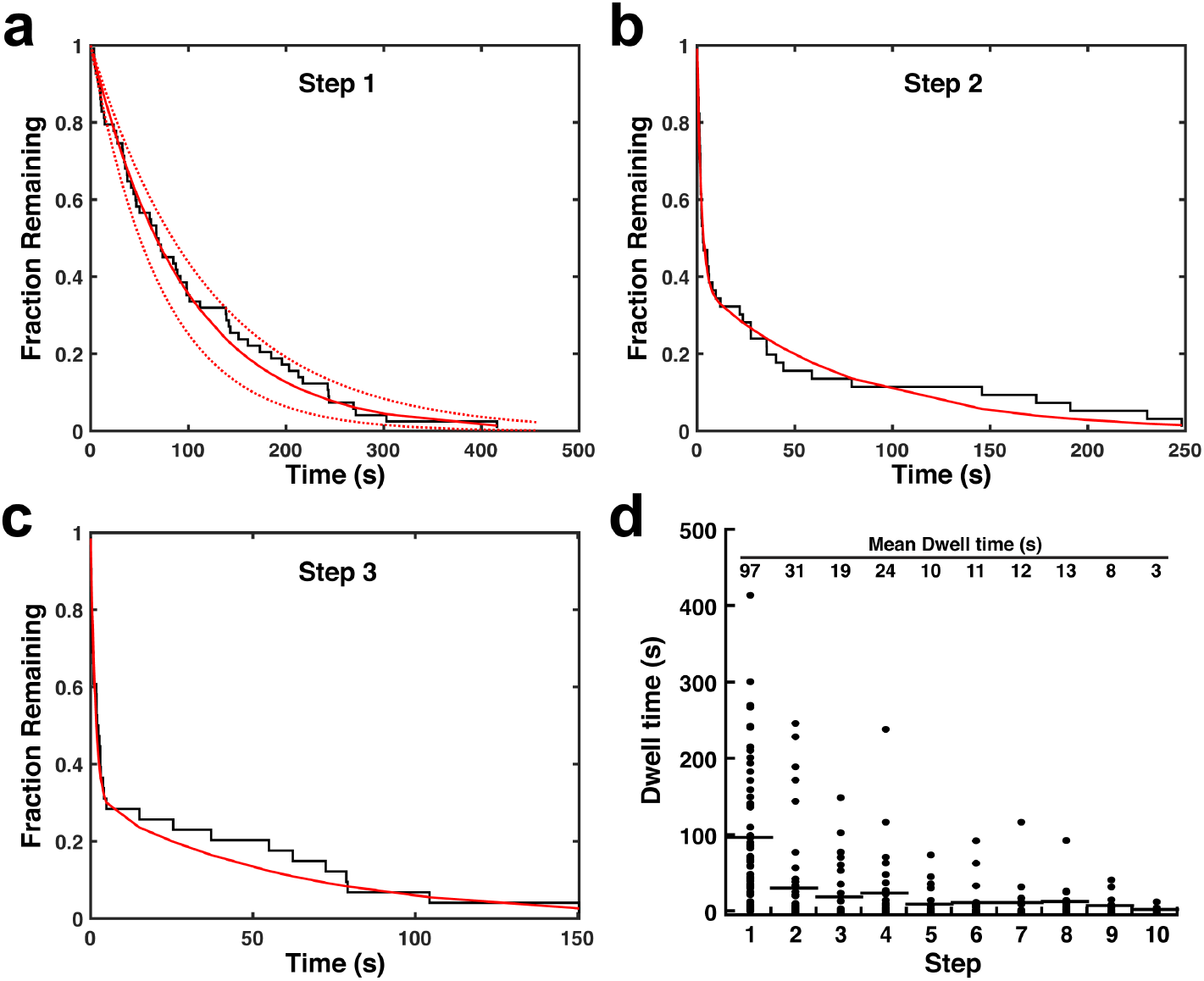
Dissociation from the anchor site controls product release by telomerase. **a-c**, Inverse cumulative distribution of dwell times for the **a** first, **b** second, and **c** third step of all telomerase trajectories (black staircase, n = 60 traces). Solid red line indicates a single exponential fit, dashed lines the 90% confidence intervals. **d**, Dot plot of individual dwell times for the first 10 steps of all telomerase trajectories (n = 60). Mean dwell time is indicated at the top of the graph and by the black horizontal lines.

## Discussion

Telomere maintenance is essential for the survival of stem cell populations and the vast majority of cancer cells. Telomerase is thought to elongate telomeres in a single processive elongation event, but how telomerase remains bound to the chromosome end through repeated cycles of RNA-template translocation was a key unanswered question. The singlemolecule analysis of processive telomerase catalysis presented here provides direct evidence that telomerase tightly associates with substrate DNA at an anchor site, allowing multiple telomeric repeats to be synthesized before the product is released (Fig. 7). In addition, we demonstrate that the product synthesized by telomerase folds into GQ structures and that GQ formation by telomeric DNA is more dynamic than previously thought (Fig. 7).

**Fig. 7.**
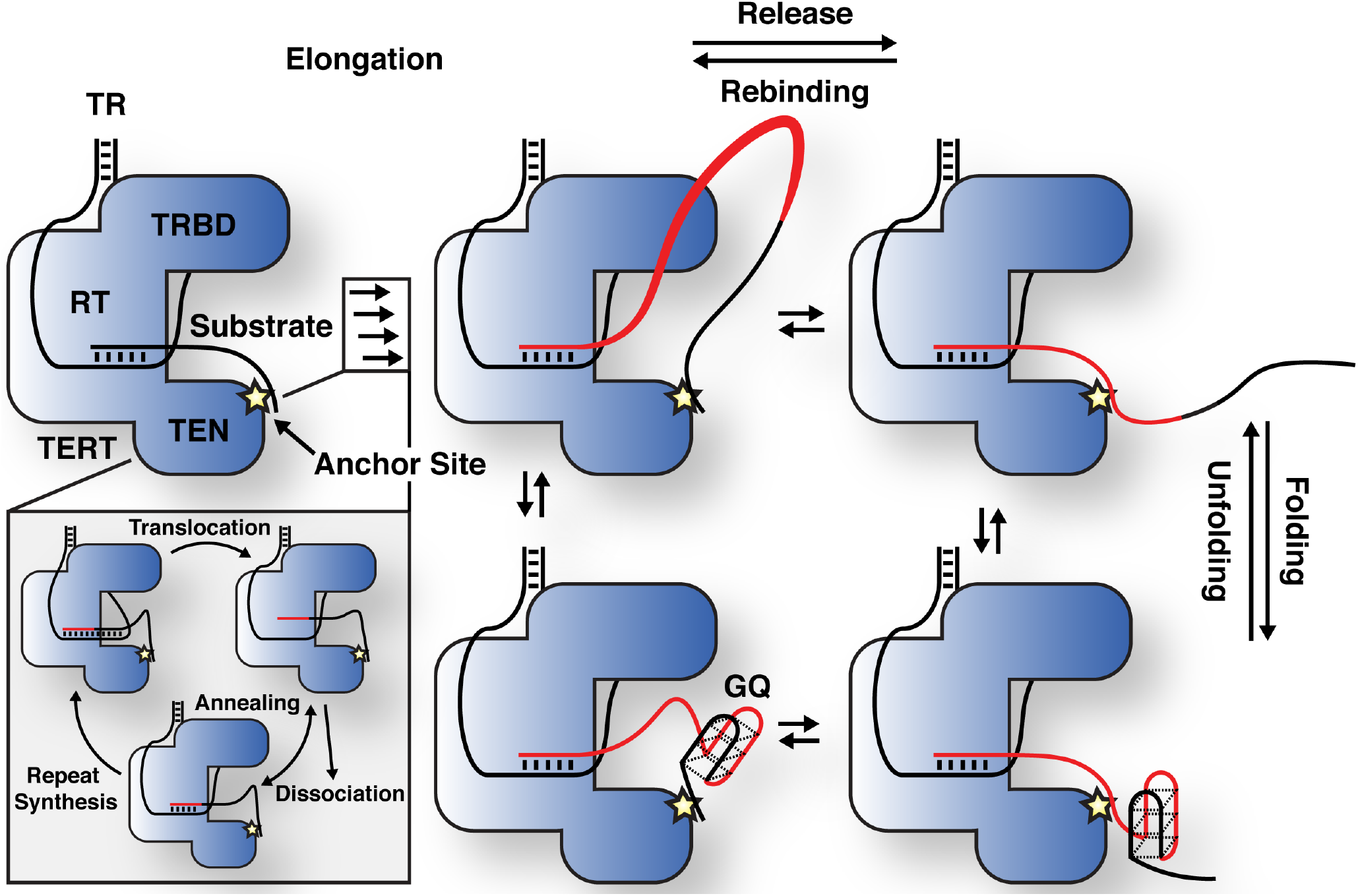
Model for processive telomerase catalysis. Telomerase associates with substrate DNA by base-pairing with the telomerase RNA template and at an anchor site. Multiple cycles of repeat synthesis, RNA-DNA melting, RNA translocation, and re-annealing occur without product being released from the anchor site, leading to the formation of a product loop. When the product DNA dissociates from the anchor site multiple telomeric repeats are released in a single large step. After and potentially before release, the product DNA can dynamically fold and unfold G-Quadruplex structures.

### An Anchor Site in Telomerase Facilitates Processive Telomere Elongation

The presence of a second DNA binding site within telomerase, in addition to RNA-DNA base pairing, was proposed soon after the identification of telomerase ^25, 39–41^. Since telomerase had been demonstrated to be processive ^7^, but RNA-DNA base pairing had to be disrupted to allow RNA template translocation, the presence of an anchor site was an attractive model to explain how telomerase can synthesize more than a single telomeric repeat without dissociating from its substrate. Since then, a number of studies have assessed the DNA binding properties of TERT from various yeast species ^28, 42^, ciliates ^27, 43, 44^, and humans ^18, 45, 46^. These reports analyzed individual TERT domains, TERT in the absence of TR, or relied on crosslinking, and therefor did not conclusively demonstrate the existence of an anchor site. The data presented here strongly suggest that the fully assembled human telomerase RNP actively engaged in catalysis contains a secondary DNA binding site. Our results demonstrate that multiple telomeric repeats are synthesized without release of the nascent DNA from telomerase. Since processive repeat synthesis by telomerase requires temporarily disrupting RNA-DNA base pairing, an additional anchor site must be present to prevent product release (Fig. 7).

### The Molecular Nature of the Telomerase Anchor Site

While our observations imply that human telomerase contains a secondary binding site that associates with the DNA substrate, this study does not provide insight into the molecular nature of this anchor site. Reports by others have implicated the TEN-domain of TERT from various species, including human, in DNA binding. The TEN-domain is positioned adjacent to the catalytic ring formed by the TRBD, RT, and CTE of TERT, placing it in close proximity to the DNA substrate ^13^, the ideal location to provide anchor site function. In telomerase from T. thermophila the TEN-domain does not directly contact the substrate ^14^. In fact, p50 and Teb1, which are homologs of human TPP1 and POT1, obstruct the TEN-domain from accessing the short DNA fragment present in the structure. Importantly, there are key differences between human and T. thermophila telomerase when it comes to DNA binding. The core RNP (TERT and TR) of T. thermophila binds to DNA with much lower affinity than human telomerase (maximal half-lives of 2 min and >20 hours, respectively) ^44, 47^, indicating that human telomerase might contain additional DNA contact points. It is therefore possible that the human TEN-domain provides anchor site function, while T. thermophila telomerase relies on its co-factors to bind to substrate DNA. Based on previous studies and its positioning in the structure of telomerase the TEN-domain is a likely candidate to harbor the anchor site, but we cannot rule out that other TERT domains or telomerase RNP components contribute to anchor site function.

Similar to the anchor site, the POT1/TPP1 complex also binds to single stranded telomeric DNA ^48^. POT1/TPP1 forms a direct interaction with the TEN-domain of TERT ^15–17^, stimulates telomerase processivity ^49^, and increases the affinity of telomerase for substrate DNA^38^. In future studies, it will be critical to determine how the DNA-binding activities of telomerase and POT1/TPP1 are coordinated to facilitate processive telomere elongation.

### The Contribution of the Anchor Site to Processive Telomerase Catalysis

Interactions between telomerase and the single-stranded overhang of chromosomes ends are so rare that they might only occur once per cell cycle ^20, 22^. But when formed, they last long enough for telomerase to compensate for the telomeric repeats lost during DNA replication in a single processive telomere elongation event ^20^. This suggests that telomerase processivity rather than the rate of nucleotide addition is the key parameter controlling the extent of telomere lengthening. Processive telomerase catalysis requires the combined action of the anchor site and repeated cycles of melting, translocation, and reannealing of substrate DNA-TR base-pairing.

Our single-molecule telomerase assay not only allowed us to demonstrate the presence of an anchor site within telomerase it also facilitated the precise quantification of its DNA binding properties. The anchor site tightly associates with substrate DNA. But, the half-life of anchor site binding (t_1/2_ = 1.1 min) is substantially shorter than the half-life of primer binding to the telomerase RNP not engaged in catalysis (minimal t_1/2_ = 15 min, maximal t_1/2_ > 1200 min) ^47^. This is consistent with the anchor site contributing to substrate binding, and RNA-DNA base pairing further increasing the affinity of telomerase for telomeric DNA when it is idle. Our measured anchor site off-rate (k_off_ = 0.01±0.001 s^−1^) in combination with a conservative estimation of the on-rate (k_on_ = 1×10^6^ M^−1^s^−1^) results in a dissociation constant of K_D_ = k_off_/k_on_ =10 nM. This dissociation constant (K_D_ = 10 nM) is comparable to the Michaelis-Menten constant for telomerase relative to the DNA substrate (K_m_ = 10-20 nM) ^50, 51^. Considering that DNA substrate turnover by telomerase is fairly slow (low kcat, K_m_ = (k_off_ + k_cat_)/k_on_), this K_m_ should be comparable to the dissociation constant of primer DNA binding to telomerase that is actively synthesizing telomeric repeats. Therefore, the close correspondence of the anchor site dissociation constant and the Michaelis-Menten constant of telomerase and substrate DNA suggests that the anchor site is the main DNA binding site when telomerase is engaged in catalysis. Consistent with this model, the half-life of substrate bound to the anchor site (t_1/2_ = 1.1 min) is almost identical to the halflife of primer bound to telomerase that is actively synthesizing telomeric repeats (t_1_/_2_ = 1.0 min) ^32^. Importantly, both the Km and half-life of primer binding to actively elongating telomerase were determined under conditions with low levels of dGTP. When dGTP is limiting, the initiation of repeat synthesis after translocation is slow (Fig. 7, inset) ^33^. Thus, telomerase is “stuck” with minimal base-pairing and the anchor site is especially important to prevent substrate dissociation (Fig. 7, inset).

The amount of DNA released in a single step from the telomerase anchor site is widely distributed (5 nt to > 100 nt, Fig. 2). This indicates that a substantial amount of single stranded DNA can accumulate tightly bound to the enzyme, potentially by forming a large product loop (Fig. 7), and suggests that anchor site function is largely uncoupled from telomeric repeat synthesis by telomerase. In total, our results demonstrate that the association of the telomerase anchor site with substrate DNA is the major driving force preventing product dissociation from telomerase and thus controls processive telomere elongation.

### The Product DNA Released from Telomerase Dynamically forms G-Quadruplexes

In addition to directly visualizing product release, we were also able to analyze conformational dynamics of the telomeric DNA synthesized by telomerase. Our results show that product DNA released from telomerase readily folds and unfolds GQ structures. Several lines of evidence support this conclusion. First, the frequency of GQ folding, which manifests itself as a reverse step (shortening in tether length), increases with tether length, because longer stretches of telomeric repeats have more opportunities to form a GQ (Fig. 3). Second, this increased frequency in tether length reductions coincides with the appearance of peaks in the step size distribution (15 nt) in both forward and reverse direction that match the predicted step size of GQ formation (Fig. 3). Third, in the presence of lithium cations, which do not support GQ formation, precisely these peaks around 15 nt in length in both forward and reverse directions are completely absent (Fig. 4). Importantly, we also observe frequent reverse steps in the presence of lithium. These reverse steps likely reflect recapture of the product DNA by the anchor site. It is possible that recapture of product DNA and GQ formation are mutually exclusive, which would increase the chances of product recapture when telomerase elongates DNA in lithium containing buffer.

GQ formation by human telomeric repeat DNA has been extensively studied using force spectroscopy and other single-molecule methods. Force-ramp measurements of human telomeric GQ structures in potassium containing solution revealed mechanical stability and unfolding/refolding force hysteresis: unfolding at >20 pN while refolding at < 5 pN under typical force-ramp conditions ^5, 6^. We were therefore surprised to observe both folding and unfolding of GQs at a constant force of 4.0-4.5 pN. To further confirm that GQ formation occurs in the product DNA released from telomerase, we analyzed the dynamics and step sizes of GQ formation by telomeric repeat DNA in the absence of telomerase. Under conditions identical to our telomerase assays, telomeric sequences capable of forming one or up to three GQs rapidly folded and unfolded with a step size distribution that largely overlapped with the length change predicted for GQ folding and observed in our telomerase trajectories (Fig. 2, Fig. 5). In addition to the close correspondence in step size, the mean dwell time of GQ folding and unfolding in the absence of telomerase (11 s) was similar to the mean dwell time prior to length changes in extended telomerase products (10 s). Together our observations strongly support our interpretation, that GQ folding and unfolding occurs in the product DNA released from telomerase.

### General Implications for Telomere Maintenance

The results presented in this study address a long-standing question: How does telomerase remain bound to its DNA substrate when RNA-DNA base pairing is disrupted during the translocation step of the telomerase catalytic cycle? We demonstrate that the anchor site within telomerase constitutes a high affinity DNA binding site, which prevents substrate dissociation during processive telomeric repeat synthesis. The anchor site facilitates synthesis of multiple telomeric repeats (2-20 repeats were observed, Fig. 2), which could loop out from the catalytic core, prior to product release in a single large step (Fig. 7). This suggests that telomerase can synthesize a sufficient amount of telomeric repeats to compensate for the 50 nucleotides lost during DNA replication without releasing the chromosome end from the anchor site. Furthermore, since telomerase can bind a large loop of single stranded DNA it is also possible that the anchor site captures the chromosome end at an internal site rather than directly at the 3’ end. Future studies will have to address, how anchor site activity is coordinated with the DNA binding activity of POT1/TPP1.

The dynamic nature of GQ formation we observed has important implications for telomere maintenance by telomerase. GQs formed by the 3’ overhang at chromosome ends inhibit the association of telomerase with its substrate. While POT1 has been shown to promote GQ unfolding ^52^, it too competes with telomerase for DNA binding. Given these circumstances, how can telomerase ever access the chromosome end? Our results suggest that GQs formed at the chromosome end could stochastically unfold, without being aided by POT1 or other factors, to allow telomerase to bind to its substrate. The recruitment of telomerase to telomeres via TPP1 would increase the local concentration of telomerase and allow it to associate with the chromosome end when it becomes available for binding. Whether GQ formation directly influences the rate of telomerase catalysis or product release, as has been suggested by others ^4^, remains to be addressed in future studies.

In total our results provide a comprehensive picture of the molecular mechanisms underlying processive telomerase catalysis, which represent an important advance in our understanding of telomere maintenance, a cellular process that plays a central role in human aging, as well as cancer formation and survival ^1^.

## ACKNOWLEDGEMENTS

We thank Jayakrishnan Nandakumar and Iain Cheeseman for comments on the manuscript. This work was support by grants from the NSF (MCB-1919439) to M.J.C. and the NIH (R00 GM120386) to J.C.S. J.C.S. is a Damon Runyon Dale F. Frey Scientist supported (in part) by the Damon Runyon Cancer Research Foundation (DFS-24-17). This preprint was generated using a template from Ricardo Henriques.

## AUTHOR CONTRIBUTIONS

Experiments were designed by E.M.P., M.J.C., and J.C.S. Optical trapping with telomerase was carried out by E.M.P. Optical trapping with multi G-quadruplex construct was carried out by E.M.P. Optical trapping with single G-quadruplex construct was carried out by J.S. and B.P. All other sample preparation and experiments were carried out by E.M.P. and J.C.S. E.M.P., M.J.C., and J.C.S. analyzed data and wrote the manuscript.

## Methods

### Telomerase expression and purification

Telomerase was expressed in HEK293T cells as previously described ^53^. 30×10^6^ cells were transfected with 15 μg of TERT plasmid (3xFLAG- or 3xFLAG-HaloTag-TERT) and 60 μg of pSUPER-hTR using 150 μl of Lipofectamine 2000 (Invitrogen) in a total volume of 7.5 ml of OPTI-MEM (Gibco). Cells were harvested and lysed with CHAPS buffer (10 mM TRIS-HCl pH 7.5, 1 mM MgCl_2_, 1 mM EGTA, 0.5% CHAPS, 10% glycerol, 5 mM beta-mercapto-ethanol) 48 hours after transfection (100×10^6^ cells). Telomerase was bound to Anti-FLAG-M2 affinity gel (Sigma-Aldrich, A2220, 100 μl of resin), washed three times with washing buffer (20 mM HEPES pH 7.9, 300 mM KCl, 2 mM MgCl_2_, 1 mM EDTA, 1 mM DTT, 1 mM PMSF, 0,1% Triton X-100, 10% glycerol), and eluted with 200 μl of wash buffer supplemented with 0.15 mg/ml 3xFLAG peptide (Sigma-Aldrich). Telomerase quantity, composition, assembly, and biotin modification was assessed with western and northern blots as previously described ^50^, using TR probes 1, 2, and 3 (IDT, Table S1), and antibodies against FLAG (Sigma-Aldrich, anti-FLAG-M2), TCAB1 (Proteintech, 14761-1-AP), dyskerin (Santa Cruz, sc-373956), and poly-HRP-streptavidin (Thermo Scientific. 21140). TR standards for northern blots were in vitro transcribed from PCR products using the HiScribe™ T7 high yield RNA synthesis kit (New England Biolabs). Band intensities were quantified using ImageQuant TL 8.2 (GE Healthcare Life Sciences).

### Direct telomerase primer elongation assay

Telomerase activity assays were carried out for 1 hour at 30 °C as previously described in reaction buffer (50 mM TRIS-HCl pH 8.0, 50 mM or 150 mM KCl, 1 mM MgCl_2_) ^54^. Assays with telomerase alone were carried out with 50 nM primer A, 50 mM KCl, 10 μM or 50 μM dATP, dTTP, and dGTP, and 0.33 μM dGTP [α-32P] (Perkin Elmer). Telomerase assays to assess POT1/TPP1 stimulation were carried out with 50 nM primer A5, 150 mM KCl, 100 nM POT1/TPP1, 500 μM dATP, dTTP, 3.3 μM dGTP, and 0.33 μM dGTP [α-32P] (Perkin Elmer). POT1/TPP1 was purified as previously described ^15^. Telomerase products were separated on 12% poly-acrylamide, 1xTBE, 7 M Urea, sequencing gels and detected using storage phosphor screens (GE Healthcare Life Sciences, BAS-IP MS 3543) and an Amersham Typhoon IP (GE Healthcare Life Sciences). Phosphorylated primer A-2 was used as a loading control. Activity was quantified as total radioactive incorporated per lane normalized to the loading control. To compare activity between 3xFLAG- and 3xFLAG-HaloTag containing telomerase, specific activity was calculated by dividing the activity measured by telomerase assays by the TR levels determined by northern blot. Processivity was quantified as the intensity of products longer than 6 repeats divided by the total signal per lane.

### Optical trapping measurements

All optical trapping measurements were carried out at 23 °C. Optical trapping was carried out using a home-built timeshared dual optical trap combined with confocal single molecule fluorescence microscope instrument as previously described in detail ^36, 55, 56^. Briefly, an acousto-optic modulator (AOM) deflected a 1,064 nm laser (IPG) between two positions to create two independent optical traps (66.7 kHz modulation rate). A PID (proportional-integral-derivative) feedback loop stabilized the trap laser intensity to maintain constant trap stiffness (0.2-0.3 pN/nm) during trap positioning. Bead positions were measured at 66.7 kHz sampling rate using the transmitted trap laser light by infrared-enhanced quadrant photodiodes (Pacific Silicon Devices) via the standard back focal plane interferometry method. Bead position and trap stiffness calibrations were performed via standard fitting of the bead Brownian motion power spectra. A 532 nm fluorescence excitation laser was coaligned between the dual traps at the center of the tether and interlaced with a second AOM to be on only during trap laser off intervals. Fluorescence emission was collected confocally and split by color into standard donor (Cy3) and acceptor (Cy5) emission channels and detected by a pair of single photon counting avalanche photodiodes (APD). Instrument control and data acquisition were performed using a field programmable gate array-based PC card system (National Instruments PCIe-7852R). Instrument control software was written in LabVIEW version 2012 (National Instruments).

The substrate handle (SH) was generated by annealing the substrate handle oligo-nucleotides 1 and 2 (IDT, Table S1). The biotin-digoxigenin handle (BDH) was generate by PCR using biotin and digoxygenin modified oligos 1 and 2 (IDT, Table S1) and pBR322 as template. To establish tethers, the telomerase-substrate complex was formed by incubating 3xFLAG-HaloTag(biotin)-TERT containing telomerase with the SH for 30 minutes at room temperature, prior to binding of the complex to 1 μM diameter polystyrene beads (Spherotech) functionalized with an anti-digoxigenin antibody (Roche) for 10 minutes at room temperature. The BDH was incubated with a 50-fold excess of neutravidin (Thermo Scientific) for 10 minutes at room temperature, prior to binding to anti-digoxigenin antibody modified polystyrene beads ^57^. Tethers were formed by holding the beads close together in telomerase reaction buffer lacking nucleotides (50 mM TRIS-HCl pH 8.0, 50 mM KCl, 1 mM MgCl_2_, 1% glucose, 1 mg/ml glucose oxidase, 0.13 mg/ml catalase, 1 mg/ml Trolox) under continuous laminar flow in a multi-channel home-made chamber ^58^ to establish the linkage between the neutravidin on the BDH and the biotin-modified 3xFLAG-HaloTag-TERT telomerase. Stable tethers were subjected to force-extension measurements at a ramp rate of 100 nm s^−1^, which were fit to worm-like chain model DNA polymer modelling parameters. Measurements of telomerase catalysis were initiated in blank buffer. After a control period of approximately 30 seconds, tethers were transferred into reaction buffer supplemented with 50 μM dATP, dTTP, and dGTP. To determine the contribution of G-quadruplex folding and unfolding on product extension by telomerase, trapping experiments were carried out in buffers containing 50 mM LiCl instead of KCl.

To analyze GQ folding and unfolding G-quadruplex forming oligos GGG(TTAGGG)_3_ and (TTAGGG)_12_ were purchased from IDT (Table S1). GQ constructs were generated by ligating the oligos to DNA handles amplified from Lambda DNA and digested with TspRI (right handle, RH) and pBR322 digested with HindIII or PspGI to make the left handle (LH). The RH and LH were amplified using primers modified with digoxygenin and biotin, respectively (Table S1). FRET donor and acceptor fluorophores flanking the single GQ sequence were added by annealing and ligating Cy3 and Cy5 labeled 9 nt long oligos to complimentary sequences flanking the GQ sequence. The GQ construct was incubated with the streptavidin coated beads (Spherotech) for 1 hour and tethers were formed in situ by establishing a digoxigenin-anti-digoxigenin linkage of the RH with anti-digoxygenin antibody coated beads. Stable tethers were first subjected to force extension at 100 nm s^−1^ spanning a force range from 0-30 pN to unfold any pre-existing structures and to assess tether quality. The trap distance was then set to apply a constant force of 4-4.5 pN and measurements were carried in force-feedback mode.

Template elongation and GQ-dynamics were monitored as a change extension in the tether length by tracking the bead position within the movable trap. The optical trap was operated under active force feedback, where bead positions within the traps were monitored and kept constant by adjusting the location of one trap, thereby applying a constant force of 4-4.5 pN to the tether. The recorded motion of the trap tracked the changing extension of the tether e.g., the elongation activity of telomerase, release and rebinding of product, folding and unfolding of GQ structures etc. The change in location of the trap was calibrated to the equivalent change in the number of tether ssDNA nucleotides (nt, e.g., due to the catalytic elongation of the tethered DNA substrate) via standard ssDNA polymer modeling (via the extensible freely jointed chain model, persistence length = 0.75 nm, contour length = 0.56 nm/nt, stretch modulus = 800 pN). Force extension were carried out at 100 nm s^−1^. The models for tether extension vs force are produced by the sum of the standard polymer models of dsDNA representing the handles (extensible worm-like chain model, persistence length = 53 nm, contour length = 0.34 nm/bp, stretch modulus = 1200 pN) and the relevant quantity of ssDNA ^59–61^.

### Data Analysis

Offline data analysis was performed using standard methods implemented via custom codes programmed in MATLAB version 2018b (Mathworks). Data collected at 66.7 kHz was boxcar averaged to a final lower rate, as specified, but generally sufficiently fast to accurately determine telomerase template elongation states and lifetimes, between 1.5 and 24 ms per data point. Automatic state and step finding in telomerase trajectories was carried out using the code XL-ICON which is an implementation of the infinite Hidden Markov Modelling (iHMM) method which is a Bayesian nonparametric extension of the very commonly used HMM method and allows for state discovery with simultaneous correction for high-resolution tweezers measurement drift and finite response time ^62^. Regions of a small number of traces (4 out of 60) were excluded from contributing to the state and step size distributions because they had a very large number of reversing states that would have obscured the contributions from the remainder of traces. The overall extension rate was calculated by dividing the maximal extension of each trace by the time point at which it was first observed. Step finding for GQ trajectories was carried out in a simpler manner using standard HMM methods because the total length changes were limited to the folded and unfolded extensions unlike the highly processive telomerase elongation data. All trajectories were manually inspected for accuracy of the automated state and step finding. Step sizes in nm were converted to nucleotides using the force dependent extension based on the freely jointed chain model using the same parameters as above. For dwell time analysis, fitting was never performed on binned data. For dwell time distributions well fit by single exponentials, the mean dwell time, which is the maximum likelihood estimator for the time constant, is reported along with the standard error of the mean to estimate uncertainty. For dwell time distributions that were well fit by the sum of two or three exponentials (e.g., the unfolding of the single GQ tether and the distribution of all multi-GQ state dwell times), fitting was performed by minimizing the least squares error in the dwell time cumulative distribution. Error was estimated by finding fit parameter one sigma (68%) confidence bounds by standard bootstrapping of the fitting using resampled data with replacement.

### Data Availability

All data generated or analyzed during this study are included in this manuscript (and its supplemental information). Reagents and raw data are available from J.C.S. and M.J.C. upon request.

### Computer Code

Custom MATLAB code used to analyze the data generated in this study is available from M.J.C. upon request.

---

**Supplementary Figure 1.**
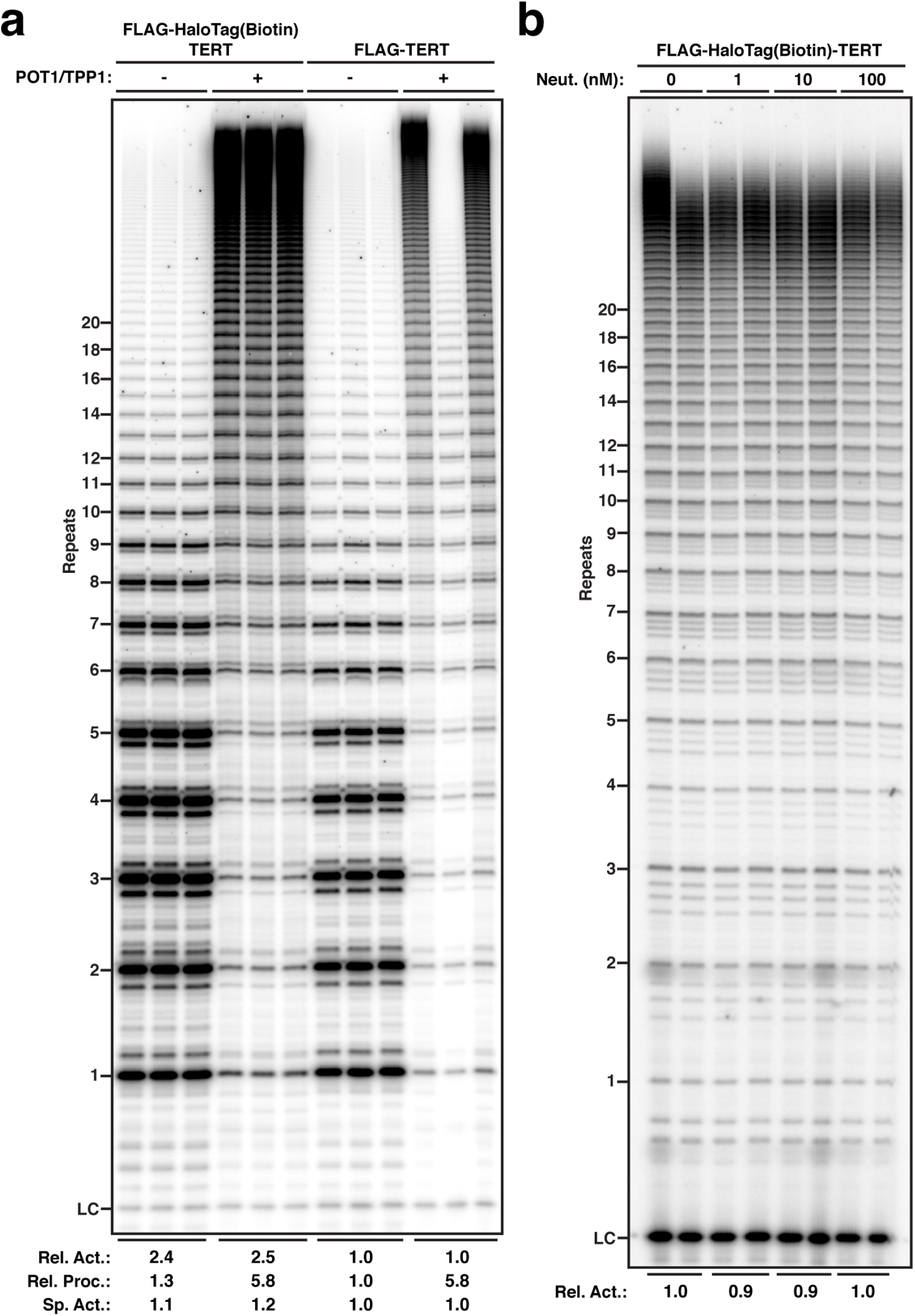
Telomerase containing a 3xFLAG-HaloTag on TERT modified with biotin is stimulated by POT1/TPP1 and its activity is unaffected by neutravidin. **a**, Direct telomerase primer extension assay of 3xFLAG-HaloTag-TERT modified with biotin and 3xFLAG-TERT containing telomerase in the absence and presence of POT1/TPP1 in buffer containing 150 mM KCl. Relative activity is total lane intensity normalized to LC and the respective 3xFLAG-TERT telomerase activity. Specific activity is additionally normalized to the relative amount of TR (Fig. 1d). Processivity is calculated by dividing the intensity of products greater than 6 repeats by total lane intensity. **b**, Direct telomerase primer extension assay of 3xFLAG-HaloTag-TERT modified with biotin in reaction buffer containing 10 μM dNTPs, 50 mM KCl, and increasing concentrations of neutravidin. Relative activity is total lane intensity normalized to LC and activity in the absence of neutravidin.

**Supplementary Figure 2.**
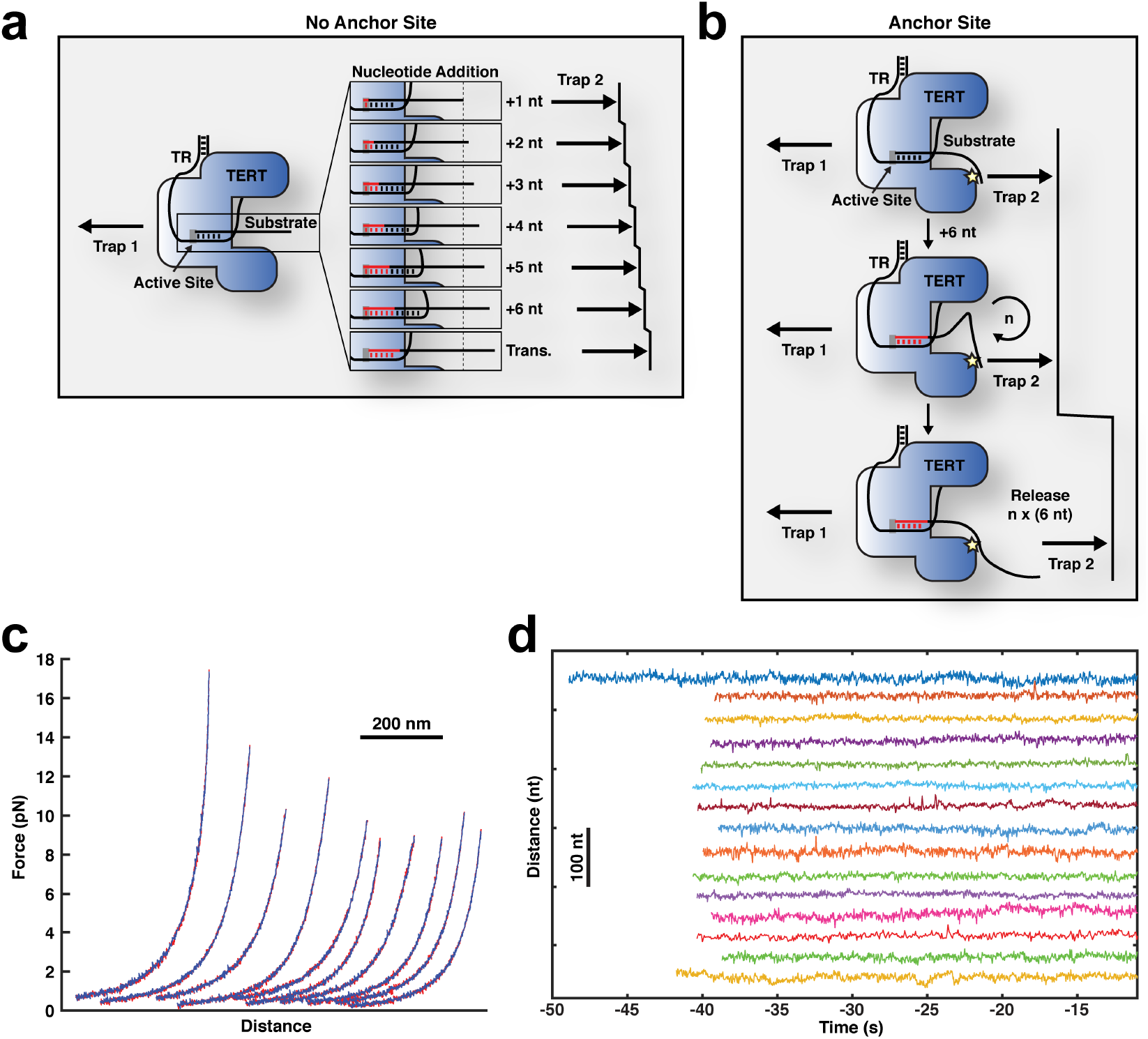
Telomerase tethers are stable in the absence of nucleotides. **a**, Expected elongation step pattern of telomerase without anchor site function. **b**, Expected elongation step pattern of telomerase with anchor site function. **c**, Force-extension curves of tethers formed by telomerase and substrate showing measurements (red) and polymer model (blue). **d**, Representative time trajectories (30 ms per data point) of telomerase catalysis prior to transfer into dNTP-containing buffer.

**Supplementary Figure 3.**
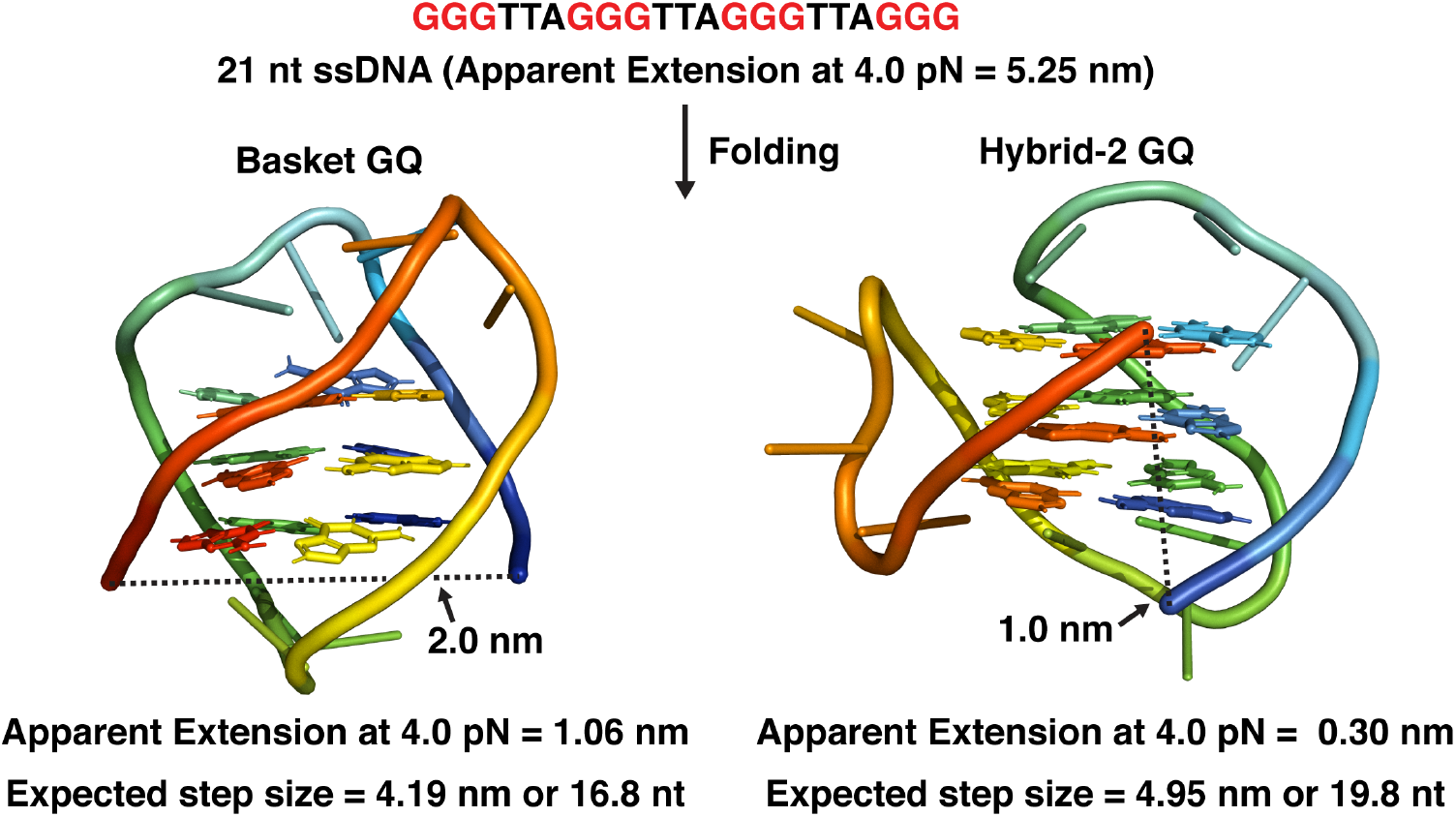
Expected extension change for the formation of a G-quadruplex structures formed by four TTAGGG repeats. Models of basket (PDB: 143D ^63^) and hybrid-2 type GQs (PDB: 2JPZ ^64^). Apparent extensions and expected step sizes were derived from the freely jointed chain polymer model.

**Supplementary Figure 4.**
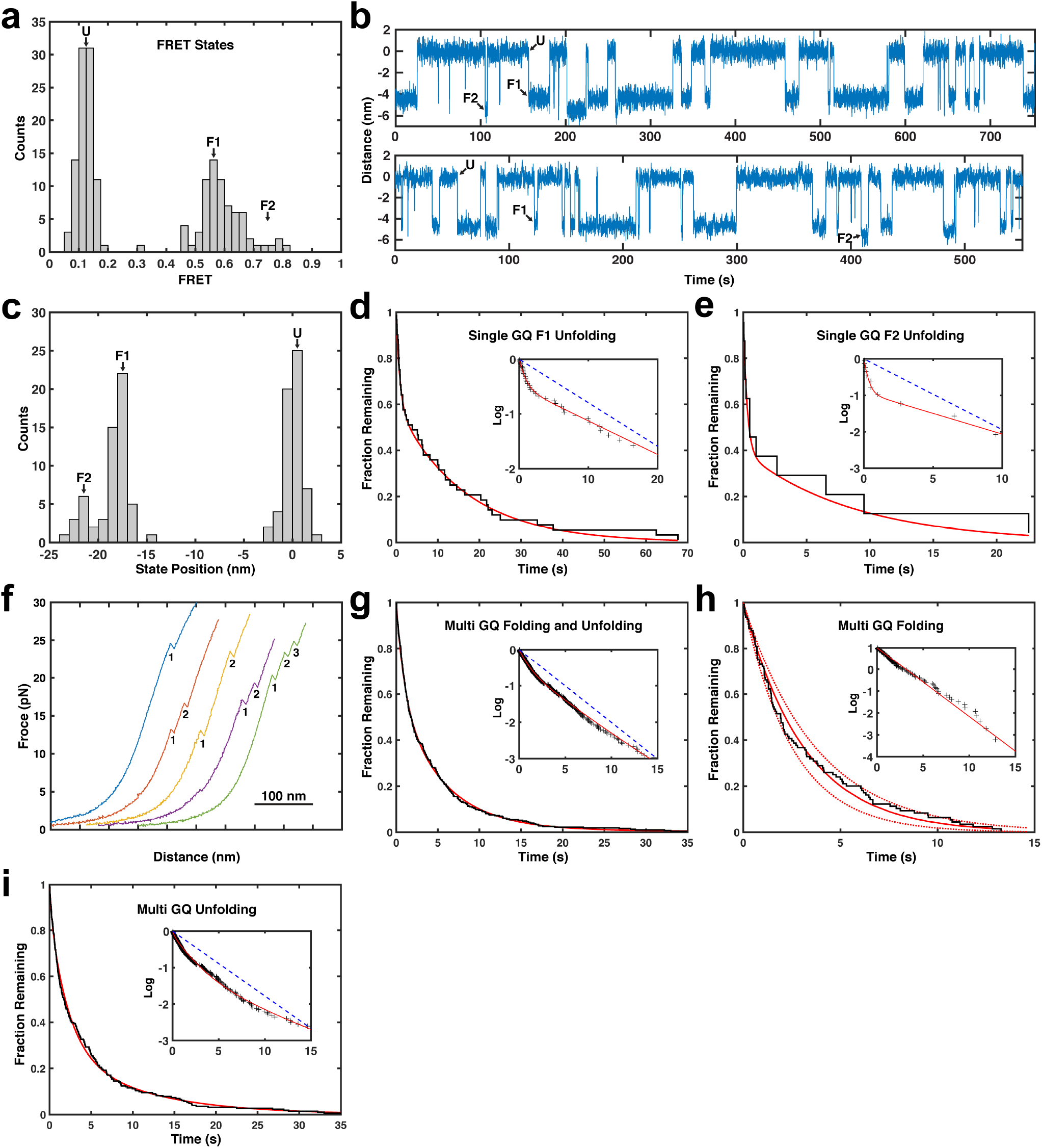
Analysis of G-quadruplex formation using high-resolution optical tweezers. **a**, Distribution of FRET states from single GQ tethers (n = 10 tethers, 163 states). **b**, Traces of two long-lasting tethers used to analyze unfolding rates of folded states F1 and F2 (72 ms per data point). **c**, Distribution of trap position states for two long-lasting tethers used to analyze unfolding rates of folded states F1 and F2 (n = 2 tethers, 117 states). **d**, Inverse cumulative distribution of dwell times for unfolding steps from folded state F1 of single GQ tethers (black staircase, n = 2 tethers, 46 dwell times). Solid red line indicates fit to the sum of two exponential decays (*τ*_u1_=0.86±0.56 and *τ*_u2_=16.0±5.2, weights of 40±14% and 60±13%). Inset: Same distribution and fit on LOG scale, blue dashed line indicates single exponential decay fit. **e**, Inverse cumulative distribution of dwell times for unfolding steps from folded state F2 of single GQ tethers (black staircase, n = 2 tethers, 12 dwell times). Solid red line indicates fit to the sum of two exponential decays (*τ*_u1_=0.29±0.4 and *τ*_u2_=8.8±6.5, weights of 60±21% and 40±28%). Inset: Same distribution and fit on LOG scale, blue dashed line indicates single exponential decay fit. **f**, Force extension curves of (TTAGGG)_12_ tethers. Numbers mark unfolding events. **g**, Inverse cumulative distribution of dwell times for folding and unfolding steps of multi GQ tethers (black staircase, n = 6 tethers, 339 dwell times). Solid red line indicates fit to the sum of three exponential decays (*τ*_u1_=1.0±0.5, *τ*_u2_=5.0±1.5, and *τ*_u3_=40±10 s, weights of 40±15%, 60±10% and 1%). Inset: Same distribution and fit on LOG scale, blue dashed line indicates single exponential decay fit. **h**, Inverse cumulative distribution of dwell times for folding steps from the completely unfolded state of multi GQ tethers (black staircase, n = 6 tethers, 102 dwell times). Solid red line indicates a single exponential fit, dashed lines the 90% confidence intervals. Inset: Same distribution and fit on LOG scale. **i**, Inverse cumulative distribution of dwell times for unfolding steps of multi GQ tethers (black staircase, n = 6 tethers, 237 dwell times). Solid red line indicates fit to the sum of three exponential decays. Inset: Same distribution and fit on LOG scale, blue dashed line indicates single exponential decay fit.

**Supplementary Figure 5.**
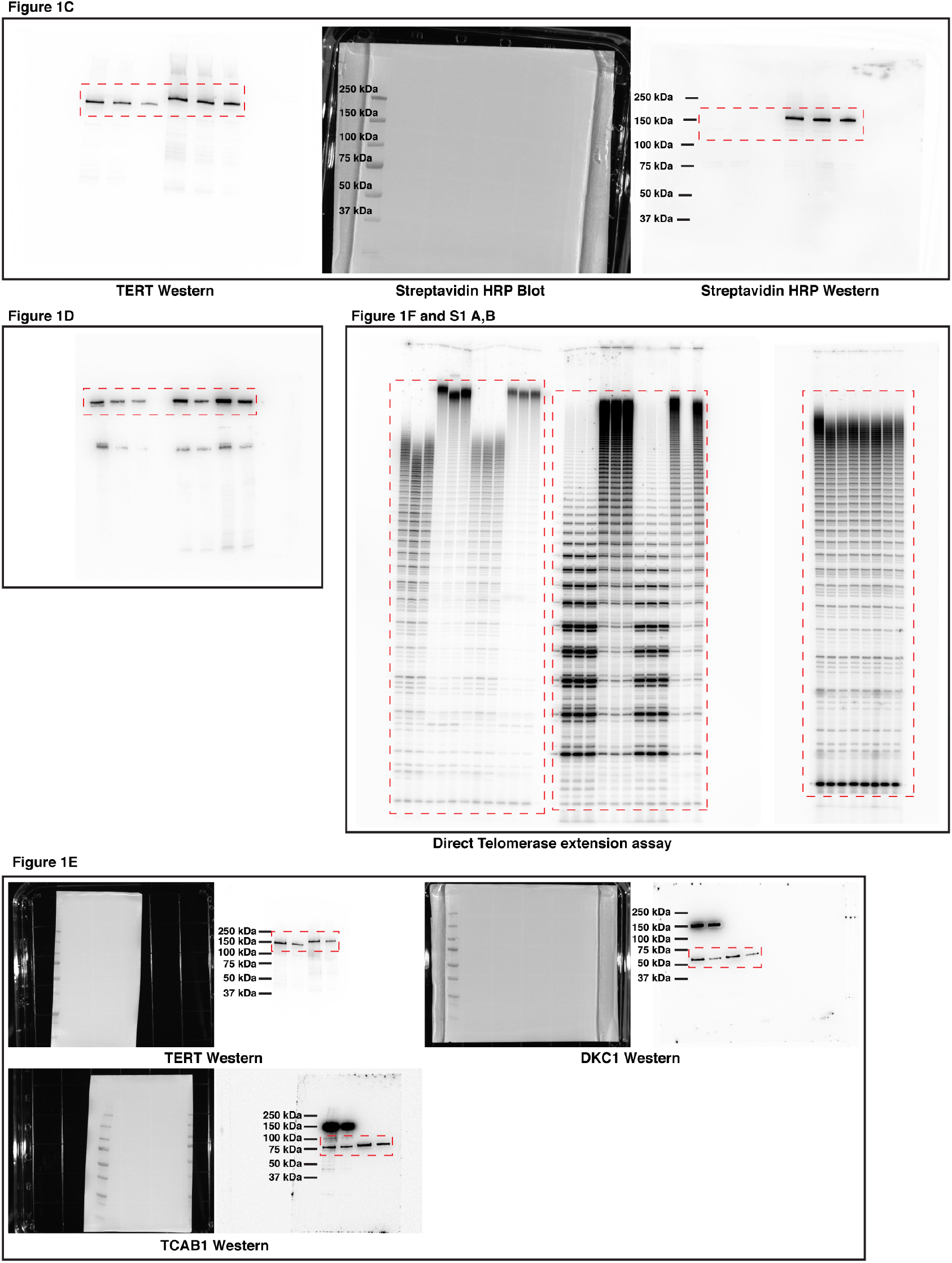
Unedited gel images for all data presented in this paper.

**Supplementary Table 1.**
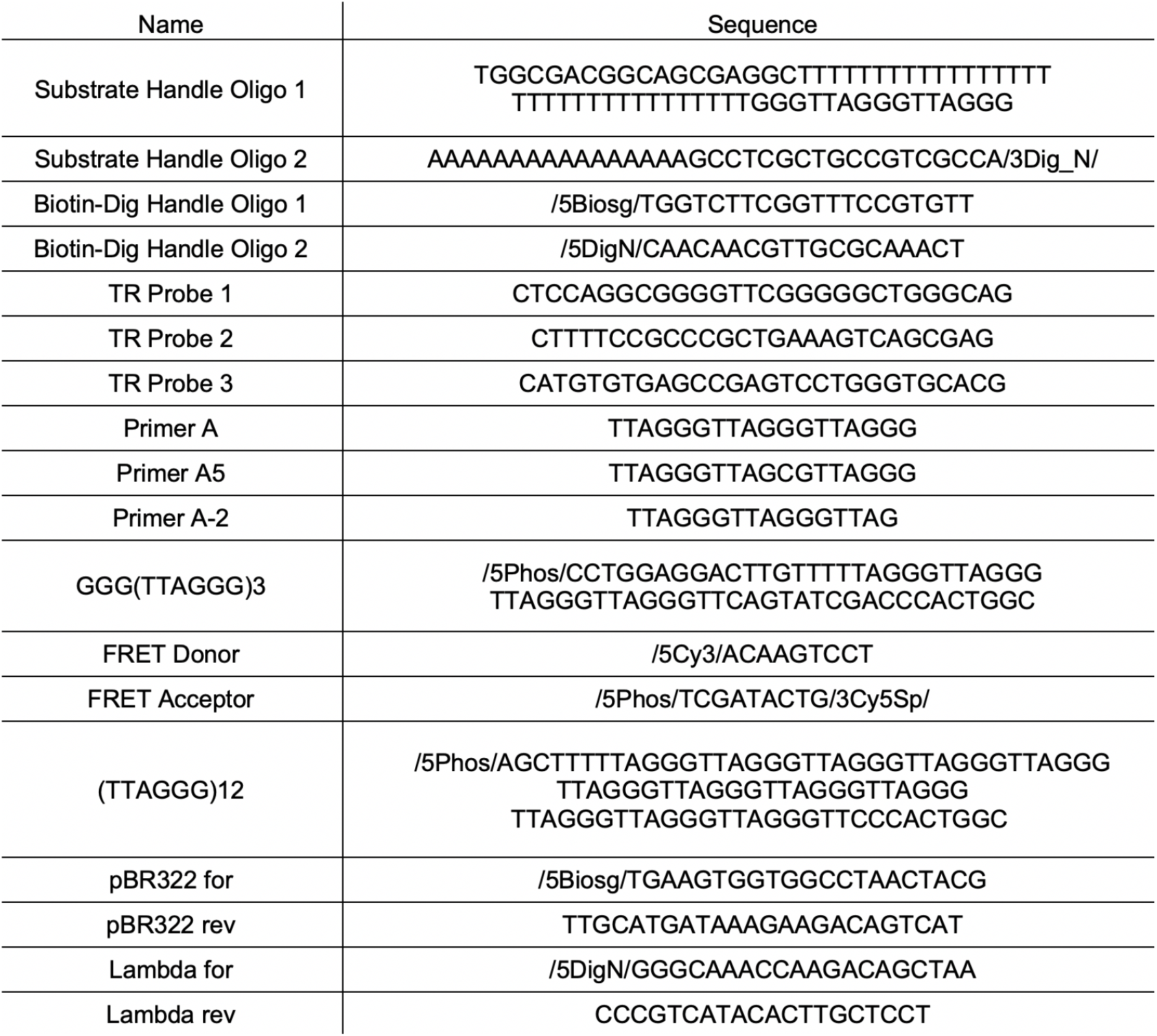
Oligonucleotides used in this study.

